# A Female Population-Averaged Musculoskeletal Model Outperforms Conventional Male-Based Generic Models in Simulating Female Gait

**DOI:** 10.64898/2026.08.23.746509

**Authors:** Ekaterina Stansfield, Hans Kainz

## Abstract

Most widely used lower-limb musculoskeletal models are derived from male anatomy and adapted to female participants solely by linear scaling, which may not capture sex-specific differences in pelvic and hip geometry. We developed a population-averaged, MRI-derived female lower-limb musculoskeletal model, built from MRI-based models of a cohort of 25 adult women using thin-plate-spline muscle-path mapping, bilateral symmetrisation, and wrapping-surface optimisation. We hypothesised that this average model, adapted to a new individual by standard linear scaling alone, would reproduce that individual’s MRI-based model’s joint kinematics, joint moments, muscle moment arms, and muscle-driven force and activation estimates more closely than a linearly scaled generic male-based model, and that this advantage would be concentrated in pelvis- and hip-dependent outputs rather than distributed evenly across all joints. Using 5-fold cross-validation, the scaled average-female model and the scaled male model were each compared against the held-out individual’s MRI-based model across gait kinematics, joint moments, muscle moment arms, muscle forces/activations, and joint reaction forces. The average-female model outperformed the male model in every output category (Holm-corrected *p* ≤ 7 ×10^−5^), supporting our primary hypothesis. Consistent with our secondary hypothesis, differences were largest and most sustained for pelvis tilt, hip flexion, and gluteal/adductor moment arms and forces, and smaller for knee and ankle kinematics. Some divergence remained localised to early-stance knee kinematics and patellofemoral loading. The population-averaged female musculoskeletal model is freely available on SimTK (https://simtk.org/projects/aver_fem) and is recommended for studies involving female participants, particularly when pelvic and hip biomechanics are the primary outcomes.

## 1. Introduction

Musculoskeletal models are widely used to estimate joint kinematics, joint moments, muscle forces, and joint contact forces that cannot be measured directly during human movement. Most widely used lower-limb models, however, are based on male anatomy and are adapted to individual participants using linear scaling of segment dimensions and masses alone (Delp et al., 1990; Rajagopal et al., 2016). This is despite well-established sex differences in pelvic and femoral morphology (White et al., 2011). These anatomical differences may influence muscle mechanics and joint loading, but their contribution is difficult to evaluate using models that do not explicitly represent female anatomy. We recently developed a semi-automatic thin-plate spline (TPS) workflow that maps homologous landmarks from a generic musculoskeletal model onto MRI-derived anatomy to generate subject-specific models (Stansfield et al., 2026). Using this approach, we showed that MRI-based personalisation alters predicted joint kinematics, joint moments, and joint contact forces even in healthy adults, and that systematic geometric differences between male and female anatomy are not captured by conventional linearly scaled generic models. However, as with other MRI- and CT-based personalisation approaches, the workflow requires imaging, registration, and muscle-path definition for every new participants, limiting its practicality for larger studies. A complementary approach was recently proposed in a Master’s thesis by Gouka (2025), who developed the MRI-based female lower-limb model YONI using the STAPLE toolbox and compared it with a scaled generic male model. The female model produced joint angles and moments that more closely matched subject-specific MRI-based models than the generic model, but the evaluation was based on a single reference anatomy and only two test participants. Previous studies have shown that incorporating personalised anatomy alters and improves musculoskeletal model predictions (Modenese et al., 2021; Kainz and Jonkers, 2023; Guggenberger et al., 2025). However, the small sample sizes of imaging-informed musculoskeletal modelling studies highlight the need for generic models that represent population-level anatomy for both sexes while retaining the simplicity of standard scaling workflows. In the present study, we developed a population-based female musculoskeletal model by averaging MRI-based models from a cohort of adult women, followed by bilateral symmetrisation and optimisation of muscle-wrapping geometry. We hypothesised that a population-based female model, fitted to new participants using conventional linear scaling, would reproduce outputs of subject-specific MRI-based models more accurately than a linearly scaled generic male model. We further hypothesised that improvements would be greatest for pelvis- and hip-related quantities, where anatomical sex differences are most pronounced.

## 2. Methods

### 2.1. Participants and MRI-based reference models

Individual MRI-based lower-limb models were available for 25 healthy adult women (age 26.2 ± 6.5 years, height 165.6 ± 7.7 cm, mass 59.6 ± 8.7 kg), generated using the TPS-based personalisation workflow described previously (Stansfield et al., 2026). We used the RajagopalLaiUhlrich 2023 model, whose bony geometry and dimensions reflect those of a 75 kg, 170 cm tall male, as a reference (Rajagopal et al., 2016; Lai et al., 2017), having adapted it to remove arms and adjust the torso inertial parameters. Personalisation was applied to the pelvis, femur, tibia, and patella; the feet and torso were left unchanged from the scaled generic male (gen-male) reference model. Briefly, the workflow maps homologous anatomical landmarks from a generic musculoskeletal model onto participant-specific MRI anatomy without requiring bone segmentation, producing participant-specific joint centres and muscle–tendon paths. Gait data (marker trajectories and ground reaction forces during walking) were collected using a 12-camera Vicon system, five Kistler force plates, and a 16-channel wireless electromyography (EMG) system (Cometa, Milan, Italy) recording eight lower-limb muscles bilaterally. Marker trajectories were acquired using an adapted Cleveland marker set (Supplementary Table S1). Participants walked at a self-selected speed along the laboratory walkway, stepping on three force plates.

### 2.2. Musculotendon and inertial parameters

Musculotendon parameters (optimal fibre length and tendon slack length) were linearly scaled according to each participant’s MRI-derived segment dimensions, consistent with standard OpenSim practice Hicks et al. (2015). Segment inertial parameters (mass, centre of mass, moments of inertia) were retained from the linearly scaled gen-male reference model rather than re-derived from MRI-based tissue volumes. Because this carries male-derived inertia into a female model, two further analyses were performed: a female-specific inertial variant was constructed by transferring de Leva (1996) female-to-male ratios for segment mass, centre-of-mass position and inertia onto the model’s own parameters, and measured inter-individual inertial variability was propagated through inverse dynamics by Monte Carlo simulation.

### 2.3. Average-model construction

An average female model was constructed from the individual MRI-based models using a seven-stage pipeline (A1–A7, Fig. 1). Because all individual models shared a common topology and coordinate-frame convention, body-local parameters (joint centres, muscle attachment and via points, and wrapping-surface parameters) were averaged arithmetically across participants (A1). Bone geometry used for visualisation was generated by averaging bone landmarks, warping the template bone surfaces, and enforcing bilateral symmetry (A2–A3). These display-only operations did not affect subsequent kinematic or kinetic analyses. Segment scale factors were incorporated into the body geometry (A4), after which bilateral reference frames, muscle paths, wrapping objects, spline functions, and skin markers were symmetrised (A5). Muscle-wrapping surfaces were optimised using a covariance matrix adaptation evolution strategy (CMA-ES; A6) to remove discontinuities in muscle moment-arm curves. Discontinuities were identified where the maximum stepwise change in moment arm exceeded 10 mm. Optimised parameters were subsequently reflected from right to left to preserve bilateral symmetry. Finally, hip abductor and adductor wrapping parameters were adjusted to match the MRI-based cohort-average moment-arm profiles (A7) by means of CMA-ES optimisation.

**Figure 1.**
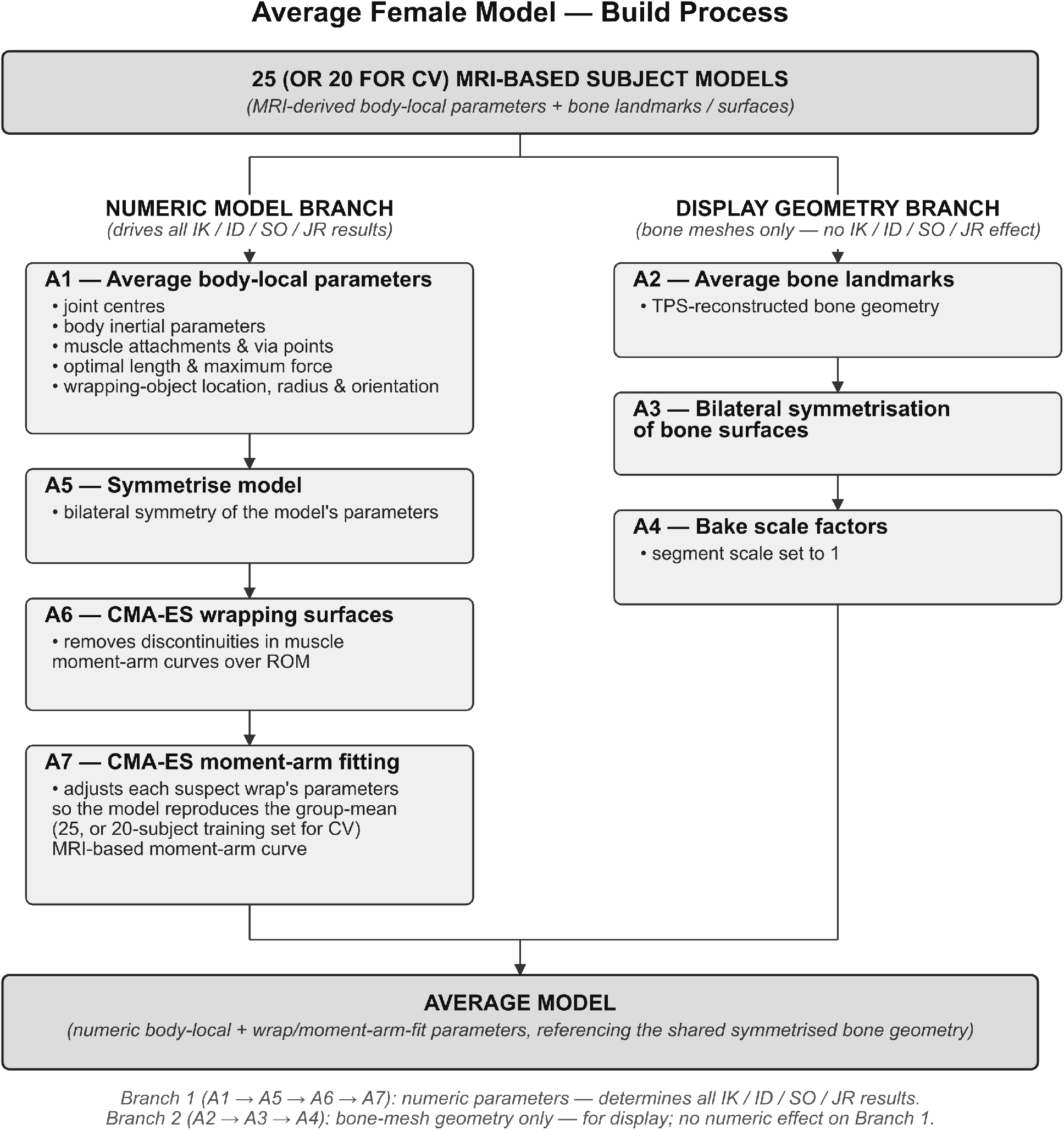
A flowchart for the process of building an average model from MRI-based personalised musculoskeletal models.

### 2.4. Cross-validation design

Model performance was evaluated using five-fold cross-validation (Fig. 2). The cohort was split into folds of five by a fixed random seed. For each fold, a new avg-female model was built from the remaining 20 individuals (training set) using steps A1, A5, A6 and A7 of the averaging pipeline, and each of the five held-out individuals was then fitted to that fold’s average model using the standard OpenSim linear (marker-based) ScaleTool (testing set). The same procedure was applied to the gen-male model (Rajagopal et al., 2016; Lai et al., 2017) as a baseline comparator, using an identical marker set and scaling configuration, so that only the underlying model geometry differed between conditions. Each held-out subject was therefore evaluated twice: once using the average female model (avg-female) and once using the gen-male model.

**Figure 2.**
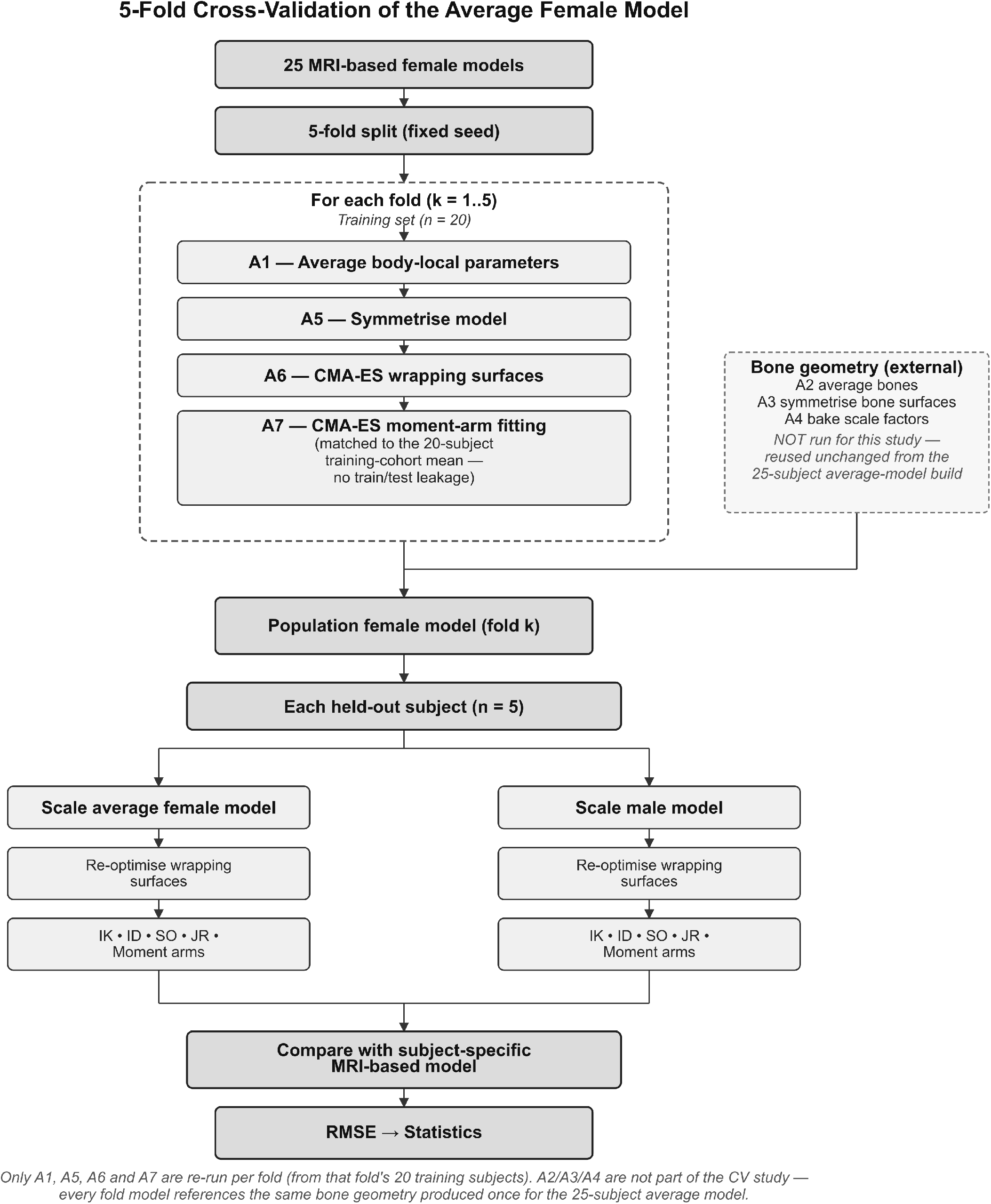
Cross-validation pipeline for testing generalisation of the average female model to unseen individuals.

### 2.5. Simulations and outcome measures

For each scaled model (avg-female, gen-male) and each participant’s MRI-based reference, joint angles (inverse kinematics), joint moments (inverse dynamics), muscle activations and muscle forces (static optimisation), and joint reaction forces were computed on one walking trial using the OpenSim 4.5.2 API. Hip abductor and adductor muscle moment arms were computed over the gait cycle for each model. For each output variable, root-mean-square error (RMSE) between the scaled model (avg-female or gen-male) and the subject’s own MRI-based reference was computed across the gait cycle.

### 2.6. Statistical analysis

For each output category (IK, ID, hip abductor and adductor moment arms, SO force, SO activation, JR), mean RMSE values were compared between the avg-female and gen-male models across all 25 held-out evaluations using paired t-tests with Holm correction for multiple comparisons. Waveform-level differences across the gait cycle were additionally assessed using paired statistical parametric mapping (SPM) t-tests (α = 0.05) for the following pairwise comparisons: gen-male vs. avg-female, avg-female vs. MRI-based, and gen-male vs. MRI-based.

### 2.7. Use of Generative AI

Generative AI assistance was used during Python code development and troubleshooting. All code was reviewed and validated by the authors, and the authors take responsibility for the resulting analyses.

## 3. Results

### 3.1. Comparing unscaled avg-female and gen-male models

Comparing geometry between the unscaled avg-female and gen-male, the male model is taller (170 cm vs. 166 cm) and heavier (75 kg vs. 60 kg). Male femur and tibia are longer, whereas some of the transverse pelvic dimensions are smaller, except for the distance between antero-superior iliac crest, i.e. ASIS, points (Table S2). At the same time, some relational measurements, such as the ratio of the lateral trochanter offset and half-hip distance, are considerably smaller in the female model.

Angle measurements, such as femoral neck-shaft angle (NSA), femur anteversion angle (FAA), and tibia torsion angle (TTA), are strongly dependent on the method of measurement Scorcelletti et al. (2020). Here, we calculated angles from bone landmarks of the gen-male and the average set of landmarks for the female cohort in the study (25 individuals), as listed in Supplementary Information. NSA and TTA are higher in the avg-female, while FAA is higher in the gen-male model. The gen-male model is not, however, representative of male anatomy in this respect: measured across our own cohorts, femoral anteversion was higher in women than in men (12.7 ± 6.2° vs. 3.6 ± 10.6°, p = 0.0006), the opposite of the model-to-model comparison (see Suppl. Information). These angle differences therefore describe the two models rather than true sex difference (Table S3).

Muscle moment arms computed over the gait cycle on the two unscaled models differed by less than 5% in eleven of 22 muscle–coordinate panels, including all ankle plantarflexors. Differences concentrated at the hip: abductor moment arms were 9.5–12.2% smaller in the avg-female model and adductor and hip-flexor moment arms 6.4–11.0% larger, following directly from its wider inter-hip distance and proportionally smaller trochanteric offset (Table S4, Figure S1).

### 3.2. Comparing geometry of the scaled avg-female and gen-male models with MRI-based gold standard

Geometric comparison of joint-centre locations and skin-marker positions relative to each subject’s MRI-based reference model showed smaller deviations for the avg-female model than for the gen-male model across nearly all joints and all 34 evaluated markers (Fig. 3).

**Figure 3.**
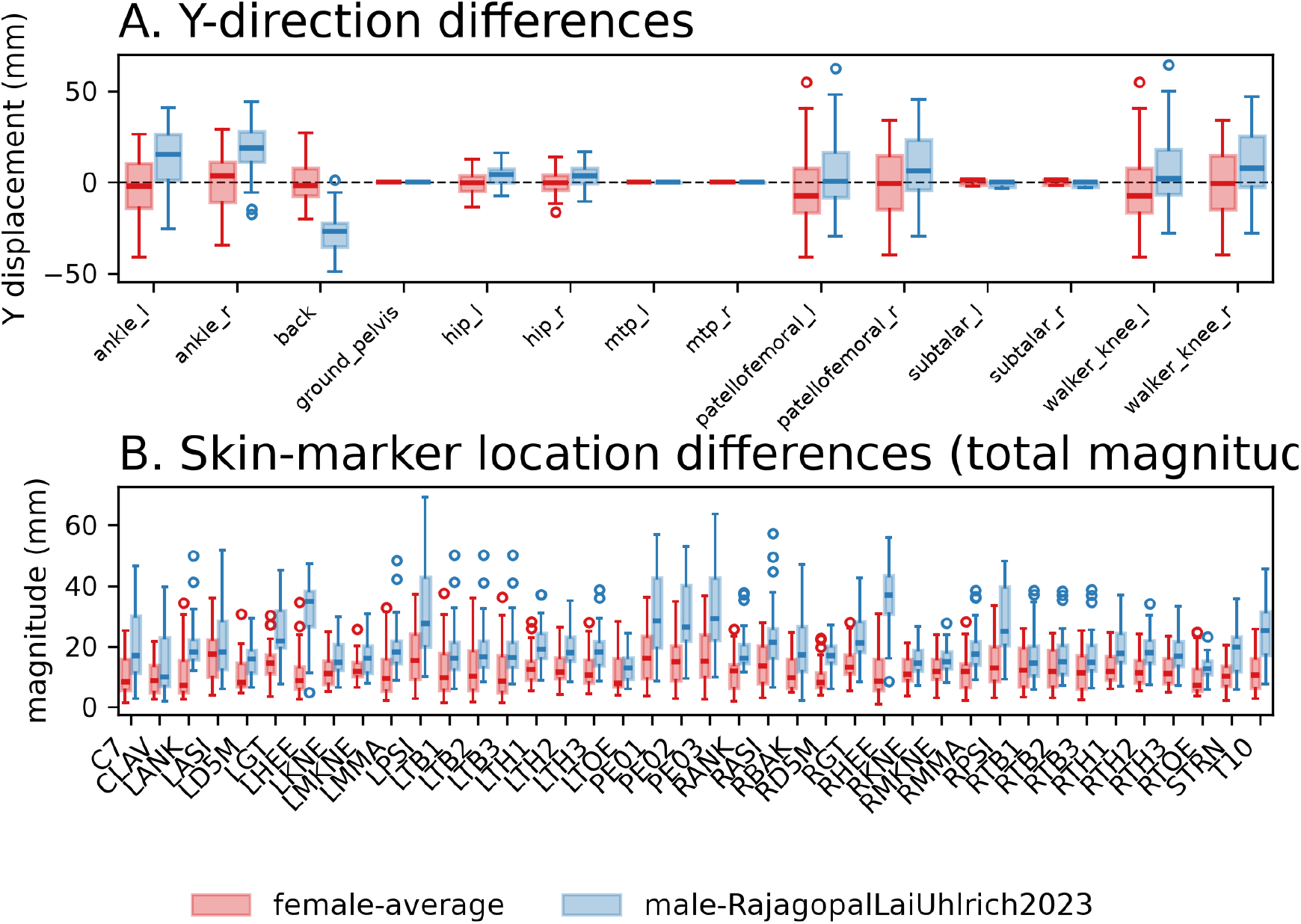
Distribution of RMSE between generic models and the MRI-based gold standard for each individual. A. Joint translation in parent frame; B. Skin marker location in the child frame.

Across all biomechanical output categories, the avg-female model showed lower RMSE compared to MRI gold-standard model than the gen-male model (Table 1). Holm-corrected paired comparisons demonstrated significant differences for all seven categories (*p* ≤ 7 × 10^−5^). The largest relative reductions in RMSE were observed for inverse kinematics and hip abduction moment arms, while smaller but significant improvements were observed for muscle forces and activations.

**Table 1.**
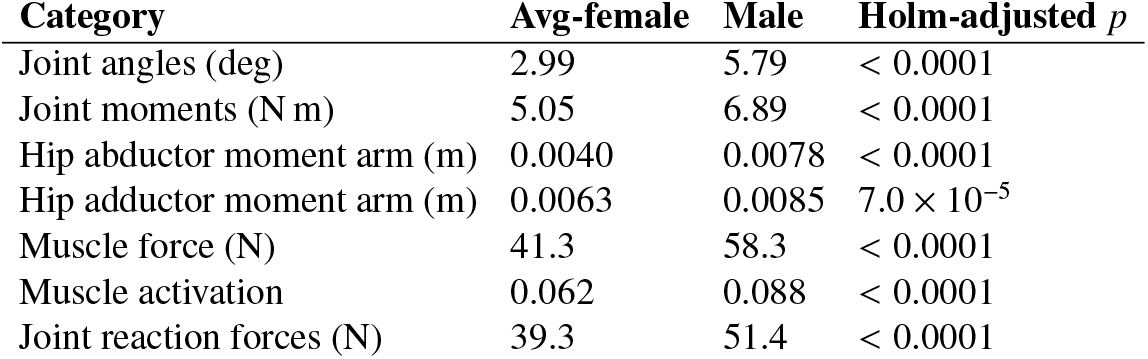
Mean RMSE by output category for the average-female and generic male models relative to each subject’s MRI-based model (*n* = 25; Holm-adjusted paired *t*-tests).

| Category | Avg-female | Male | Holm-adjusted $p$ |
| --- | --- | --- | --- |
| Joint angles (deg) | 2.99 | 5.79 | < 0.0001 |
| Joint moments (N m) | 5.05 | 6.89 | < 0.0001 |
| Hip abductor moment arm (m) | 0.0040 | 0.0078 | < 0.0001 |
| Hip adductor moment arm (m) | 0.0063 | 0.0085 | $7.0 \times 10^{-5}$ |
| Muscle force (N) | 41.3 | 58.3 | < 0.0001 |
| Muscle activation | 0.062 | 0.088 | < 0.0001 |
| Joint reaction forces (N) | 39.3 | 51.4 | < 0.0001 |

Waveform-level statistical parametric mapping (SPM) analyses showed that differences between the gen-male model and both female models (avg-female and MRI-based) were greatest for pelvis- and hip-related outputs. Pelvic tilt differed between the gen-male model and both female models throughout the gait cycle (gen-male: approximately 6–10° anterior tilt; models based on avg-female and MRI: approximately −3 to −7°), while no significant differences were observed between models based on avg-female and MRI (Fig. 4). A similar pattern was observed for ankle angle, hip flexion and hip adduction angles during early stance and swing phases (Fig. 4), hip flexion and hip adduction moments throughout the stance phase, lumbar extension moment (Suppl. Information Fig. S2), and the moment arms and muscle forces of the gluteal, iliopsoas, and adductor muscle groups (Fig. 5). In contrast, knee kinematics of the gen-male model differed from the MRI-based gold-standard model but not from the avg-female model, whereas subtalar kinematic differences between any of the three models occurred only over short portions of the gait cycle (Fig. 4).

**Figure 4.**
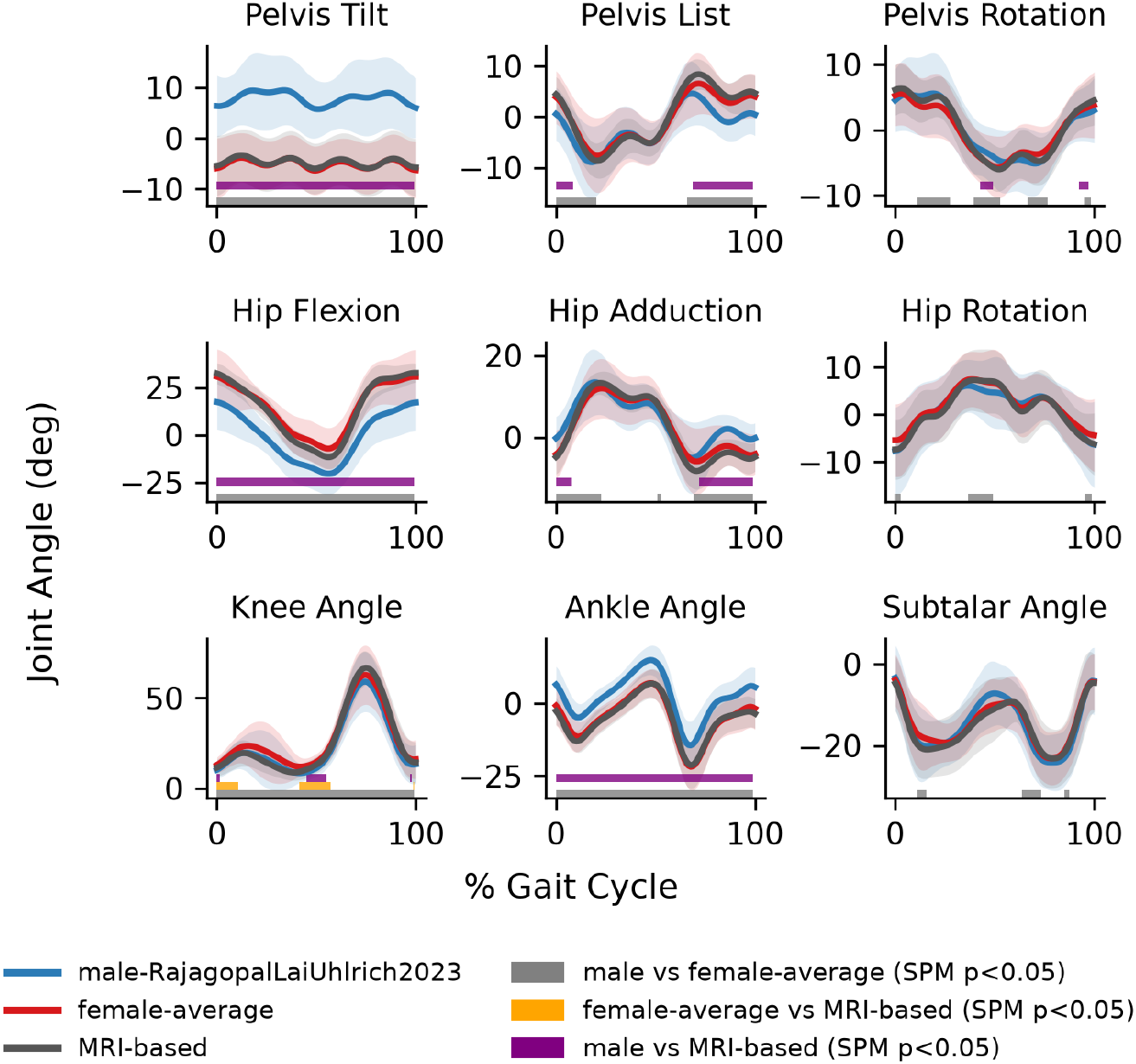
Waveforms for joint angles (Inverse Kinematics) for the right gait cycle.

**Figure 5.**
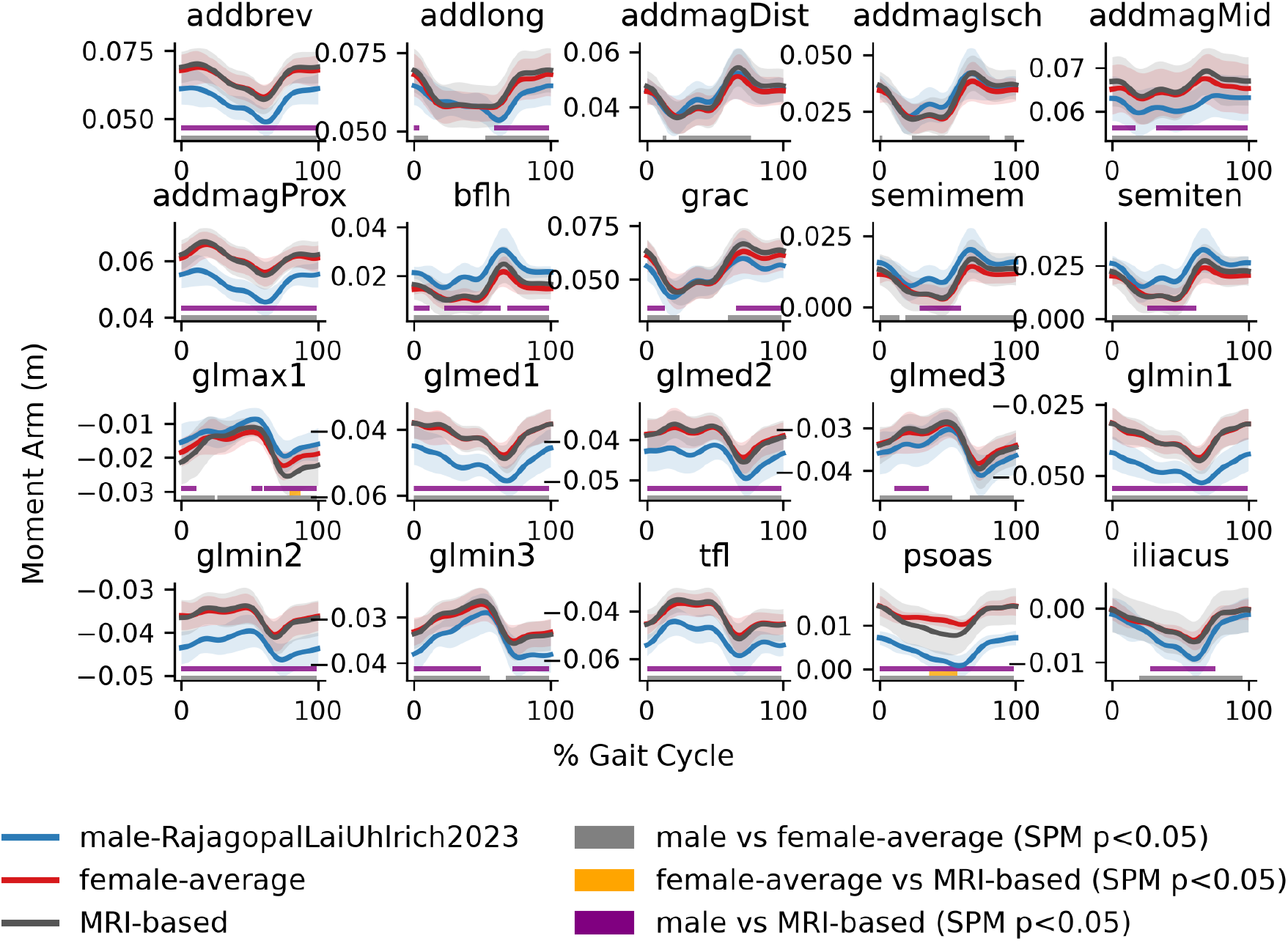
Muscle moment arms for right hip adduction degree of freedom.

**Figure 6.**
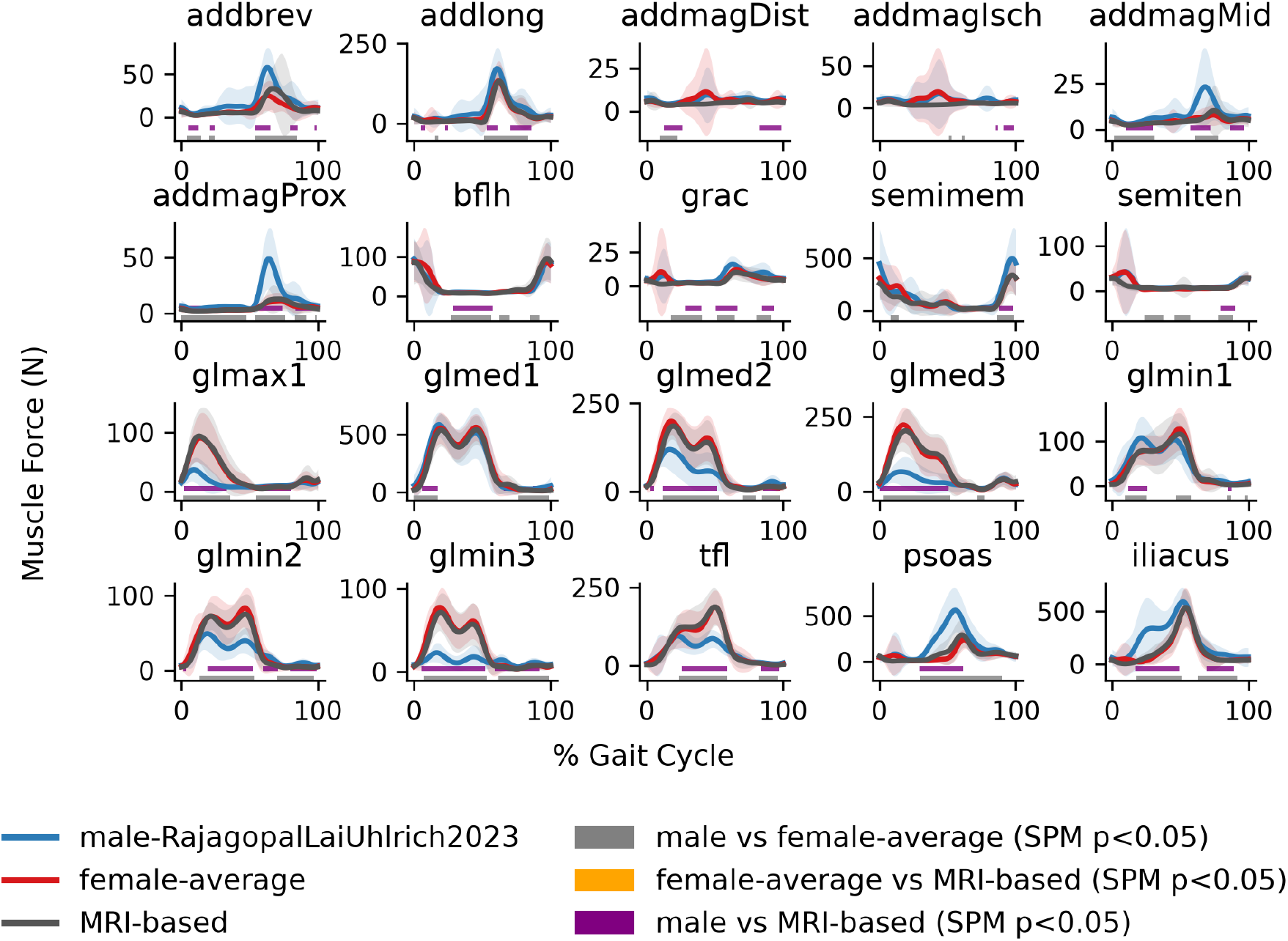
Muscle forces for selected hip ab- and adductor muscles.

Differences between the avg-female model and the MRI-based gold-standard model were localised and occurred only for selected outputs. Significant differences were detected during early- and mid-stance knee kinematics (Fig.4), patellofemoral-related joint loading (Suppl. Information Fig. S3), and a small number of individual muscle outputs, including psoas muscle moment arms (Fig.5), muscle activations in biceps femoris short head, lateral gastrocnemius, and vasti muscles (Suppl. Information Fig. S3). These differences were confined to brief intervals of the gait cycle.

### 3.3. Influence of sex-specific inertial parameters

Substituting female-specific segment inertial parameters into the avg-female model left the peak stance-phase hip and knee joint reaction forces unchanged (+0.05% and −0.06%, both non-significant); the only appreciable effect was at the lumbar joint (−8.3%, p < 0.001), driven by the torso mass multiplier (Suppl. Information Tables S5-S8, Figure S5). Monte Carlo propagation of measured inter-individual inertial variability (Durkin and Dowling, 2003) gave uncertainties of 2.1-2.6% on peak knee and hip-flexion moments, larger than the systematic male-to-female difference at those same outputs, indicating that which average inertial description the model carries matters less than the fact that it carries an average one (Suppl. Information Tables S5–S8, Figure S5).

### 3.4. Model availability

Two variants of the generic avg-female model are made available at https://simtk.org/projects/aver_fem. One is the model with male-specific inertial parameters that was used for comparisons with the gen-male and MRI-based gold standard in this manuscript, and the second one is the model with female-specific segment inertial parameters after de Leva (1996) as described above.

## 4. Discussion

Scaled to individual dimensions, avg-female model produced lower errors than the gen-male model across all evaluated biomechanical outputs in five-fold cross-validation. Improvements were greatest for joint kinematics and hip muscle moment arms, and smaller, although still significant, for joint moments, muscle forces and activations, and joint reaction forces. This pattern reflects the different dependence of these outputs on model geometry. Kinematics and moment arms are directly determined by skeletal geometry and muscle paths; therefore, anatomical differences between gen-male and avg-female models are expected to have their largest effect on these quantities (Kainz et al., 2016). In contrast, kinetic outputs are partly constrained by measured external forces (such as ground reaction forces, GRF) and by redundancy within the musculoskeletal system, which can reduce the propagation of local geometric differences into individual muscle forces or joint reaction forces. In particular, joint moments calculated by inverse dynamics mainly depend on the location of joint centres and the GRFs. Hence, they are less affected compared to moment arms, which depend on joint centres and muscle lines of action.

The strongest differences between the gen-male and avg-female models were concentrated in pelvis- and hip-dependent outputs. This is consistent with established sex differences in pelvic morphology and femoral geometry (White et al., 2011), which directly influence hip joint-centre location and muscle moment arms. In contrast, all knee-joint related outputs and muscles that cross ankle joint showed smaller differences, suggesting that linear scaling of segment dimensions captures a greater proportion of the relevant anatomical variation at these joints. These findings support the use of sex-specific geometry where anatomical differences are expected to have the greatest biomechanical consequences.

Differences between the avg-female model and each subject’s own MRI-based gold-standard were substantially smaller than differences between the gen-male and avg-female models (Fig. 3), but they were not absent. Deviations were localised primarily to knee kinematics, patellofemoral-related loading, and selected individual muscle outputs. Such differences are expected because population averaging removes subject-specific features that cannot be recovered through linear scaling alone. The purpose of the average female model is therefore not to replace full MRI-based personalisation, but to provide a more anatomically appropriate starting point for large-scale studies where individual imaging is impractical.

Our findings extend previous efforts to incorporate female-specific anatomy into musculoskeletal models. Gouka (2025) showed that an MRI-based gold-standard female lower-limb model produced joint angles and moments closer to subject-specific MRI-based models than a scaled gen-male model, but validation was limited to two test participants. Here, we evaluated a population-based avg-female model using five-fold cross-validation across 25 MRI-based gold-standard models, with waveform-level statistical comparison and assessment of residual wrapping artefacts. Unlike subject-specific MRI personalisation (Stansfield et al., 2026), the avg-female model can be applied using conventional OpenSim marker-based scaling without requiring imaging, registration, or manual definition of muscle paths. This provides a practical route for incorporating female-specific anatomy into larger biomechanical studies.

Population-based approaches to sex-specific modelling have also been explored in other anatomical regions. For example, Roos et al. (2020) developed population-representative neck models from anthropometric data, while other studies have used sex-matched anatomical atlases or single-specimen datasets to improve model geometry. Together, these efforts indicate increasing recognition that gen-male-based models may not adequately represent female anatomy. However, a validated, population-averaged female lower-limb model that can be used within standard scaling workflows has, to our knowledge, not previously been reported.

Several limitations should be considered. First, personalisation was restricted to the pelvis, femur, tibia/fibula, and patella; the feet and torso remained based on the generic reference model. Therefore, the present results describe the effects of lower-limb sex-specific geometry rather than whole-body anatomical differences. Second, the model was derived from a relatively young adult female cohort and requires validation in populations with greater variation in age, body size, and ancestry. Third, we tested segment inertial parameters that were inherited from the scaled generic model and derived from the literature de Leva (1996), rather than obtained them from MRI-based tissue volumes. We showed that a female-specific inertial variant left peak stance-phase hip and knee joint reaction forces unaffected and altered only lumbar loading, and inter-individual inertial variability exceeded the male-to-female difference at the hip and knee (Supplementary Information). Both inertial variants are released at https://simtk.org/projects/aver_fem but neither has been validated for trunk or lumbar loading, which is the one output where the inertial description materially matters. Finally, our validation was limited to walking; additional tasks such as running, stair negotiation, and dynamic balance should be evaluated.

Taken together, these results demonstrate that a population-averaged female musculoskeletal model provides a practical alternative to the male-based generic models, improving representation of female lower-limb biomechanics without requiring subject-specific imaging. The largest benefits occur for pelvis- and hip-dependent outputs, where anatomical sex differences are most directly expressed.

## Acknowledgements

This project was funded by the Austrian Science Fund (FWF) Principal Investigator Award to ES (10.55776/P35714). During the preparation of this manuscript, the authors used Claude (Anthropic) to assist with editing for conciseness. The authors reviewed and edited all AI-assisted output and take full responsibility for the content of the manuscript.

## Supplementary Information

### Skin marker set

**Table S1.**
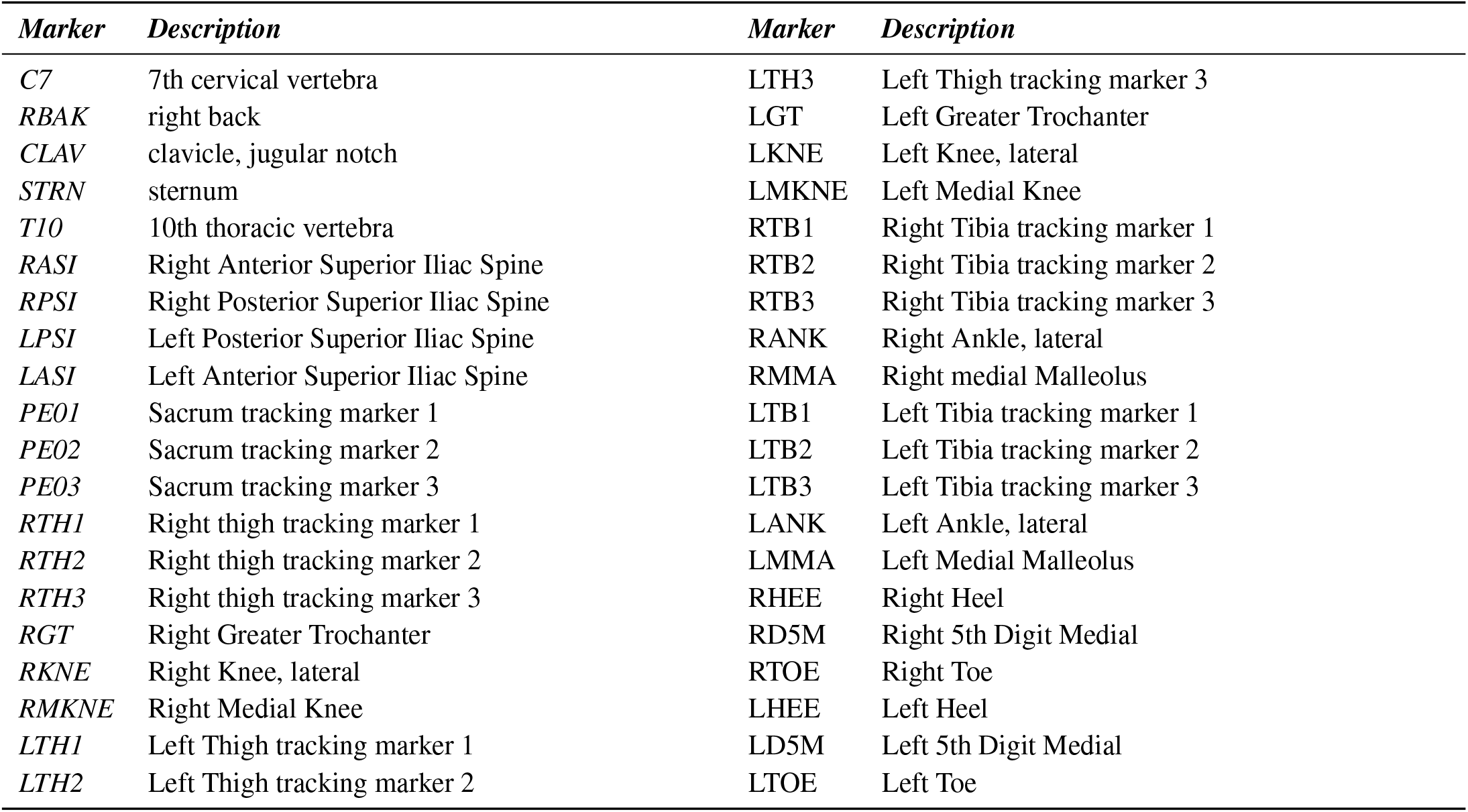
Names and descriptions of Cleveland skin markers.

| *Marker* | *Description* | *Marker* | *Description* |
| --- | --- | --- | --- |
| *C7* | 7th cervical vertebra | *LTH3* | Left Thigh tracking marker 3 |
| *RBAK* | right back | *LGT* | Left Greater Trochanter |
| *CLAV* | clavicle, jugular notch | *LKNE* | Left Knee, lateral |
| *STRN* | sternum | *LMKNE* | Left Medial Knee |
| *T10* | 10th thoracic vertebra | *RTB1* | Right Tibia tracking marker 1 |
| *RASI* | Right Anterior Superior Iliac Spine | *RTB2* | Right Tibia tracking marker 2 |
| *RPSI* | Right Posterior Superior Iliac Spine | *RTB3* | Right Tibia tracking marker 3 |
| *LPSI* | Left Posterior Superior Iliac Spine | *RANK* | Right Ankle, lateral |
| *LASI* | Left Anterior Superior Iliac Spine | *RMMA* | Right medial Malleolus |
| *PE01* | Sacrum tracking marker 1 | *LTB1* | Left Tibia tracking marker 1 |
| *PE02* | Sacrum tracking marker 2 | *LTB2* | Left Tibia tracking marker 2 |
| *PE03* | Sacrum tracking marker 3 | *LTB3* | Left Tibia tracking marker 3 |
| *RTH1* | Right thigh tracking marker 1 | *LANK* | Left Ankle, lateral |
| *RTH2* | Right thigh tracking marker 2 | *LMMA* | Left Medial Malleolus |
| *RTH3* | Right thigh tracking marker 3 | *RHEE* | Right Heel |
| *RGT* | Right Greater Trochanter | *RD5M* | Right 5th Digit Medial |
| *RKNE* | Right Knee, lateral | *RTOE* | Right Toe |
| *RMKNE* | Right Medial Knee | *LHEE* | Left Heel |
| *LTH1* | Left Thigh tracking marker 1 | *LD5M* | Left 5th Digit Medial |
| *LTH2* | Left Thigh tracking marker 2 | *LTOE* | Left Toe |

### Geometry Comparison Between the Average-Female and Generic-Male Models

For a comprehensive comparison between avg-female and gen-male models we drew data first, from the male Rajagopal-LaiUhlrich2023 (gen-male) and the average female model (avg-female), which was obtained during this project. In addition, we drew data from the original sample of 25 MRI-based models created for healthy adult women (age 26.26.5 years, height 165.67.7 cm, mass 59.68.7 kg), as well as for 22 MRI-based models created for healthy adult men (age 27.88.4 years, height 180.85.8 cm, mass 78.110.4 kg).

The average female model (avg-female) inherits the mean values for body height and body mass from the cohort of the 25 women whose MRI-based models were used for its construction. The avg-female model height is 165.6 cm and mass is 59.6 kg. In comparison, RajagopalLaiUhlrich2023 (gen-male) model is created for a man of 170 cm height and 75.3 kg body mass Rajagopal et al. (2016).

The following geometric comparisons between the two models are drawn from the landmark data collected for creating MRI-based models following Stansfield et al. (2026)(2026) and reused for the present study. Femur and tibia lengths are taken from the model joint frames, defined as the magnitude of the translation of the proximal-body offset frame of the distal joint — that is, the position of the knee joint centre expressed in the femur frame, and of the ankle joint centre in the tibia frame. The same definition is used to normalise the moment arms reported below, so the two comparisons are on a common basis (Tab. S2).

**Table S2.** Comparison of measurements between the avg-female and gen-male models.

| *Measure* | *Average-female* | *Generic-male* | *Difference* |
| --- | --- | --- | --- |
| *Femur length, joint frames (mm)* | 394.4 | 409.6 | -3.7% |
| *Tibia length, joint frames (mm)* | 395.0 | 400.1 | -1.3% |
| *Inter-ASIS distance (mm)* | 237.9 | 252.4 | -5.8% |
| *Inter-hip-centre distance (mm)* | 172.3 | 154.5 | +11.5% |
| *Inter-ischial distance (mm)* | 126.7 | 92.0 | +37.7% |
| *Pelvis depth (mm)* | 137.5 | 132.3 | +4.0% |
| *Pelvis height (mm)* | 173.6 | 195.7 | -11.3% |
| *Lateral trochanter offset (mm)* | 55.1 | 64.0 | -13.9% |
| *Trochanter offset / half inter-hip distance* | 0.64 | 0.83 | -22.8% |
| *Neck-shaft angle (°)\** | 129.8 | 120.4 | 7.8% |
| *Femoral anteversion (°)\** | 12.4 | 14.7 | -15.6% |
| *Tibial torsion, external (°)\** | 37.0 | 28.5 | 29.8% |

\* Please see the description of angle measurements below.

Angle measurements for femur and tibia are notoriously variable depending on the method. Here we derived angle measurements from the landmark data on the bones and did not expect them to match published data. In femur, the long axis was defined as a vector between the knee centre and the femur head centre, as well as its frontal plane was fit through the medial and lateral condyle points as well as through the femur head centre. The transverse plane through femur would then have the femur axis as its normal.

- Neck-shaft angle (NSA): angle, in the frontal-plane projection, between the neck axis (femur_neck_center_→ r femur_r_center) and the distal shaft direction, where the shaft axis runs from the midshaft slice centre to the slice centre 25% (proximal) of the way from the hip along the head-to-knee axis; each slice centre is the midpoint of that slice’s anterior and posterior landmarks. Measuring against the distal (rather than proximal) shaft direction yields the infero-medial angle (~ 125°) rather than its supplement.
- Femoral anteversion angle (FAA): angle, in the transverse-plane projection, between the same neck axis and the condylar axis (knee_r_lat → knee_r_med); both vectors point medially, so the resulting angle is small, with positive defined as neck-anterior-to-condylar-line.
- Tibial torsion angle (TTA): not defined in the published protocol; defined here by analogy as the transverse-plane angle between the proximal condylar axis (tibia_r_med→tibia_r_lat) and the distal malleolar axis (tibia_r_med_malleol_tip→ fibula_r_lat_malleol_tip), measured relative to a tibial long axis running from ankle centre to knee centre. Positive values denote external torsion.

NSA is higher and FAA is lower in the avg-female model than in gen-male model. Table S3 provides a cross check against our data on 25 female and 22 male participants of the study.

**Table S3.**
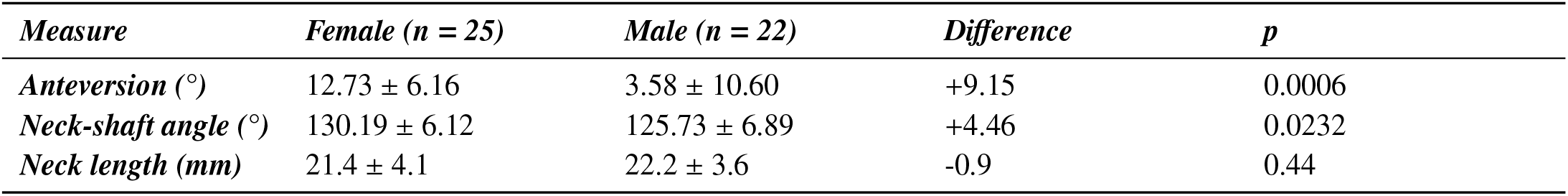
Male and female individual cohort data for femur angles.

| *Measure* | *Female (n = 25)* | *Male (n = 22)* | *Difference* | *p* |
| --- | --- | --- | --- | --- |
| *Anteversion (°)* | 12.73 ± 6.16 | 3.58 ± 10.60 | +9.15 | 0.0006 |
| *Neck-shaft angle (°)* | 130.19 ± 6.12 | 125.73 ± 6.89 | +4.46 | 0.0232 |
| *Neck length (mm)* | 21.4 ± 4.1 | 22.2 ± 3.6 | -0.9 | 0.44 |

The cross-check with avg-female and gen-male data shows that the former reproduces its own cohort’s average anatomy, while gen-male model is not representative of our male cohort. Individual male anteversion values were highly variable (SD 10.6°, with nine of 22 men returning negative values). This angle depends almost entirely on the placement of a single femoral neck landmark, so the male cohort mean should be treated with caution.

### Muscle Moment Arm Comparison Between the Average-Female and Generic-Male Models

Moment arms were computed on the unscaled models via direct OpenSim API coordinate sweeps. Sweeps covered 61 points across physiological gait ranges (hip flexion −20°̆ − 40°, hip adduction −15°̆ − 15°, knee 0°̆70°, ankle −25°̆20°) for 22 muscle-coordinate panels on the right side. Results are normalised by the relevant segment length (femur for hip/knee, tibia for ankle), with each model normalised by its own segment lengths. Both models were swept over an identical angle grid, and Table S4 reports the mean moment-arm magnitude across that range rather than a peak value, because the two models may be reaching their maxima at different joint angles.

Across the 22 muscle–coordinate panels, most normalised moment arms were similar between the two models: eleven panels differed by less than 5% in mean magnitude across the swept range, including all three ankle plantarflexor panels (soleus −0.2%, gastrocnemius medialis −0.3%) and rectus femoris at the hip (+0.4%) (Tab. S4).

The differences concentrate in three functional groups. The hip abductors were consistently smaller in the average-female model (gluteus minimus posterior −12.2%, tensor fasciae latae −11.0%, gluteus medius posterior −10.8%, gluteus medius anterior −9.5%), while the hip adductors and hip flexors were larger (adductor magnus mid +11.0%, psoas +10.4%, adductor longus +6.4%). This pattern follows directly from the pelvic geometry reported above: the female hip centres sit further apart while the trochanter lies proportionally closer to the hip centre, reducing the abductor lever arm and increasing the adductor one.

The knee extensors were smaller in the average-female model over most of the range (rectus femoris −14.8%, vastus lateralis −14.3%, vastus medialis −12.4% in mean magnitude), but this difference is strongly flexion-dependent rather than uniform: near full extension the female moment arms are marginally larger (+5%, +5.8% and +1.4% respectively at 0°), the ordering reverses between 4.8° and 6.8° of flexion, and the gap then widens progressively to approximately −33% at 70°. Tibialis anterior at the ankle was smaller by 11.3% and, unlike the knee extensors, was uniform across the whole range (−11.1% to −11.8%).

**Table S4.**
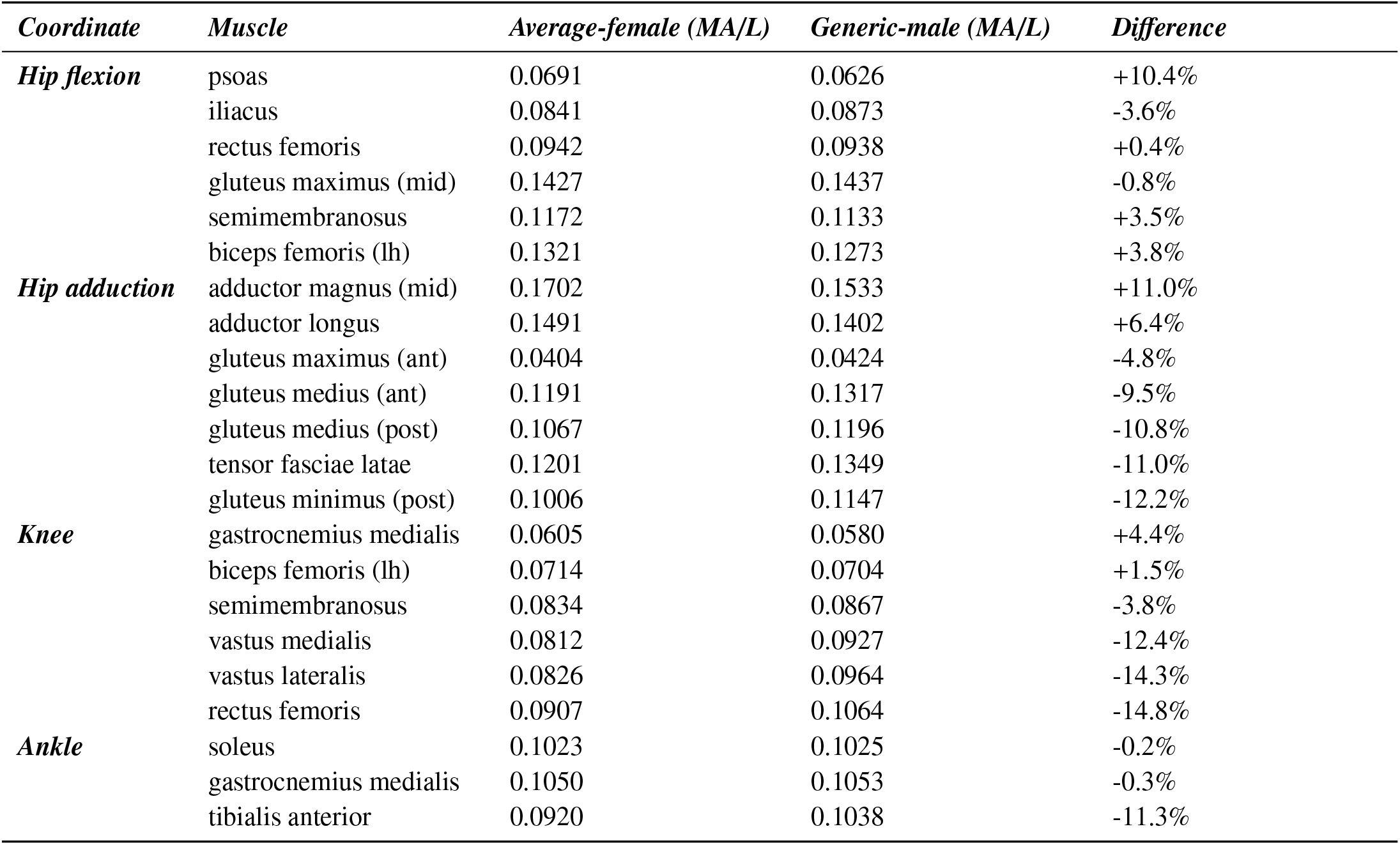
Comparison of segment-length normalised muscle moment arms. Values are dimensionless (moment arm ÷ segment length), averaged as magnitudes over the swept range; negative differences denote smaller moment arms in the average-female model.

**Figure S1.**
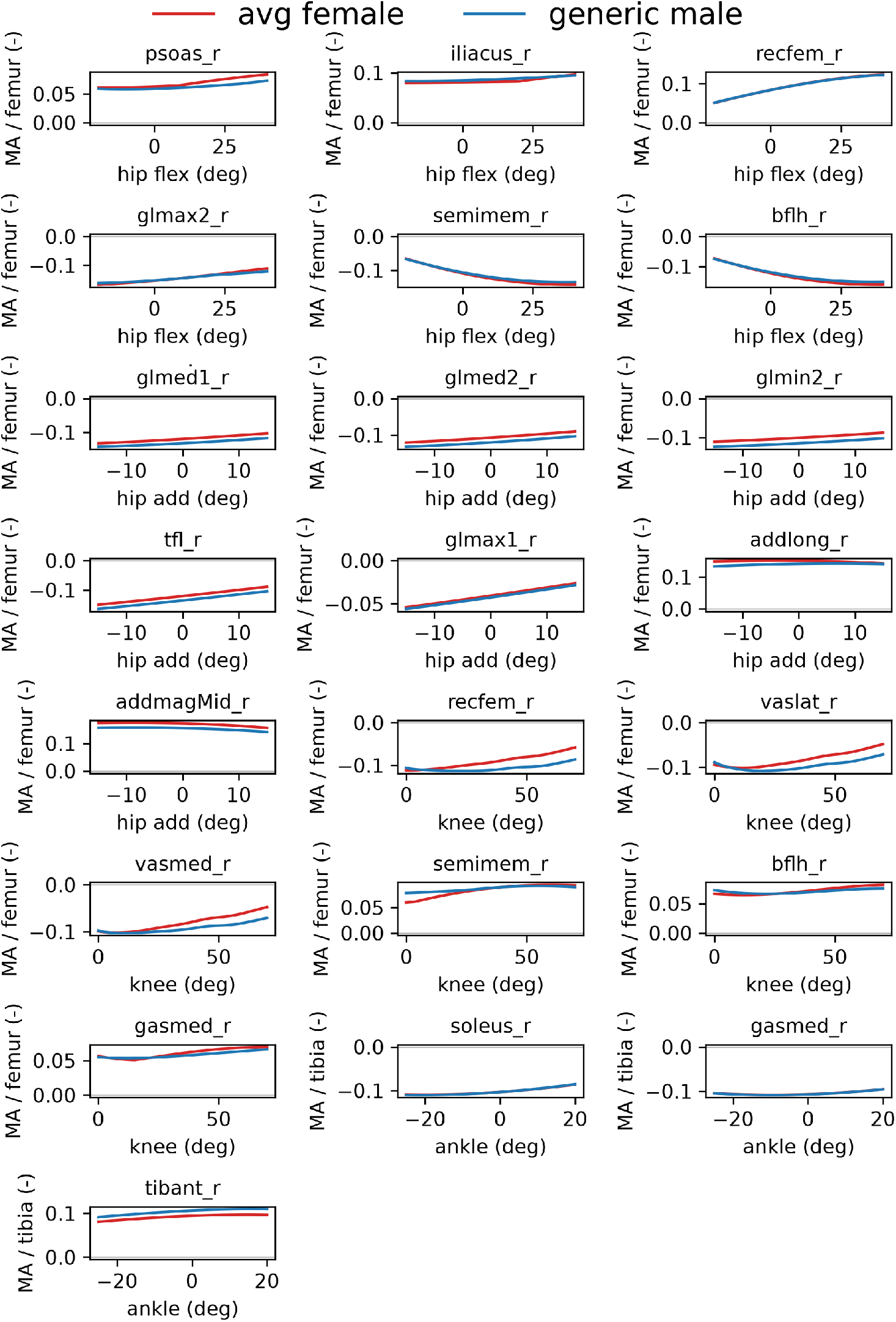
Muscle moment arms normalised by segment length, avg-female vs. gen-male.

### Comparing biomechanical outputs for scaled avg-female, gen-scaled and MRI-based gold standard models

Biomechanics results are discussed in the main manuscript.

**Figure S2.**
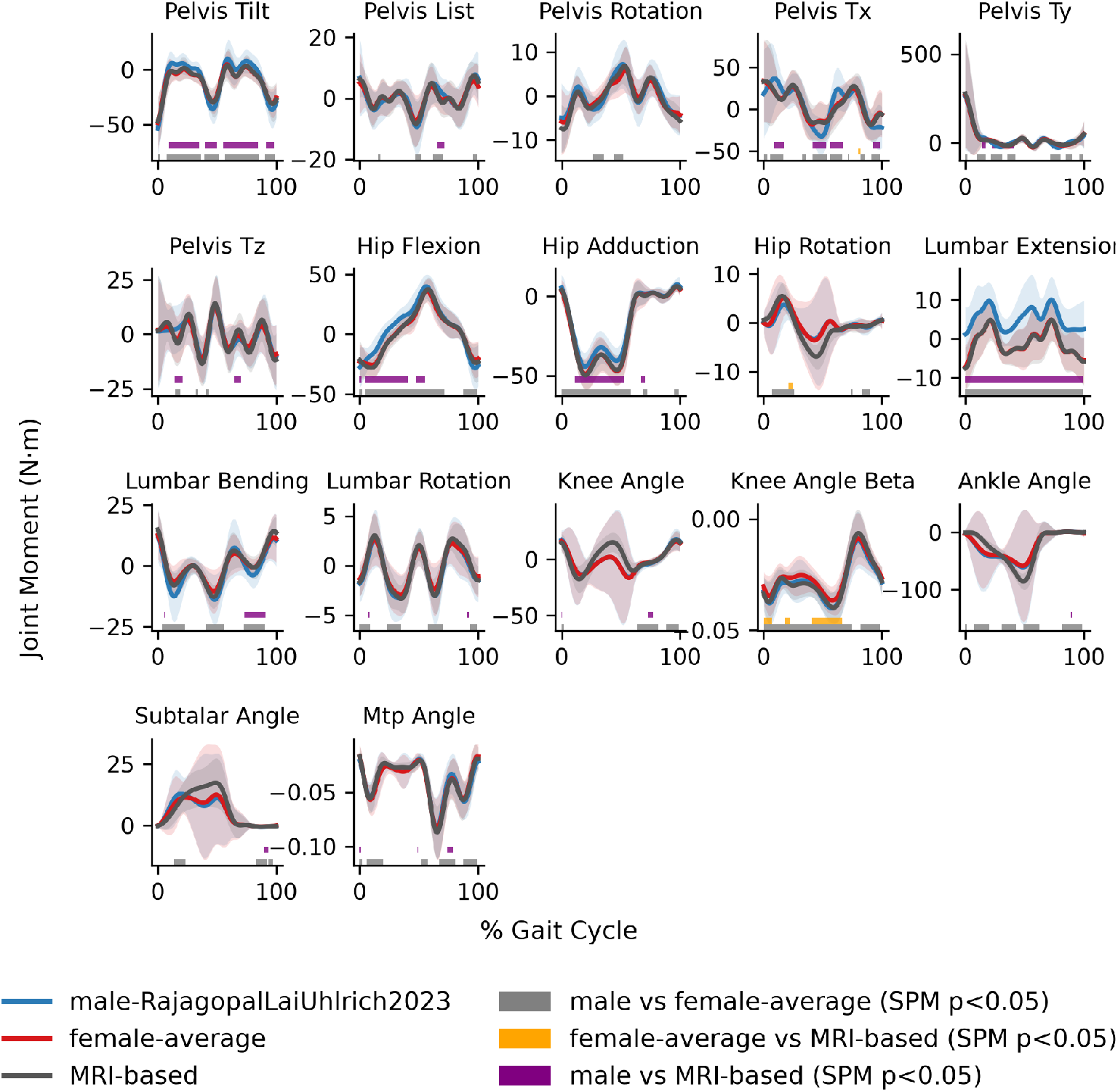
Waveforms and SPM analysis results for joint moments (Inverse Dynamics), right gait cycle.

**Figure S3.**
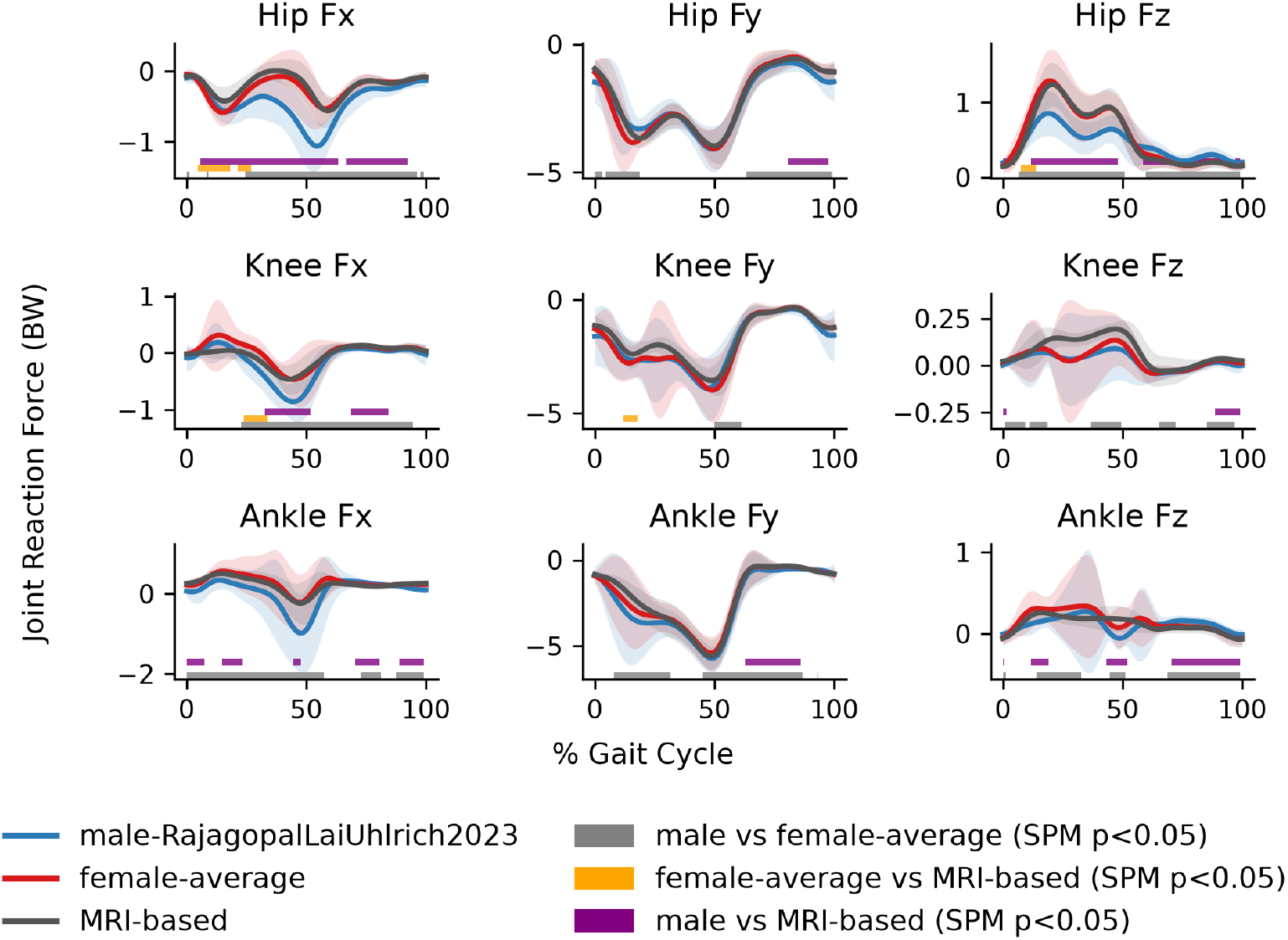
Joint reaction force waveforms and SPM analysis results for the right limb, right gait cycle.

**Figure S4.**
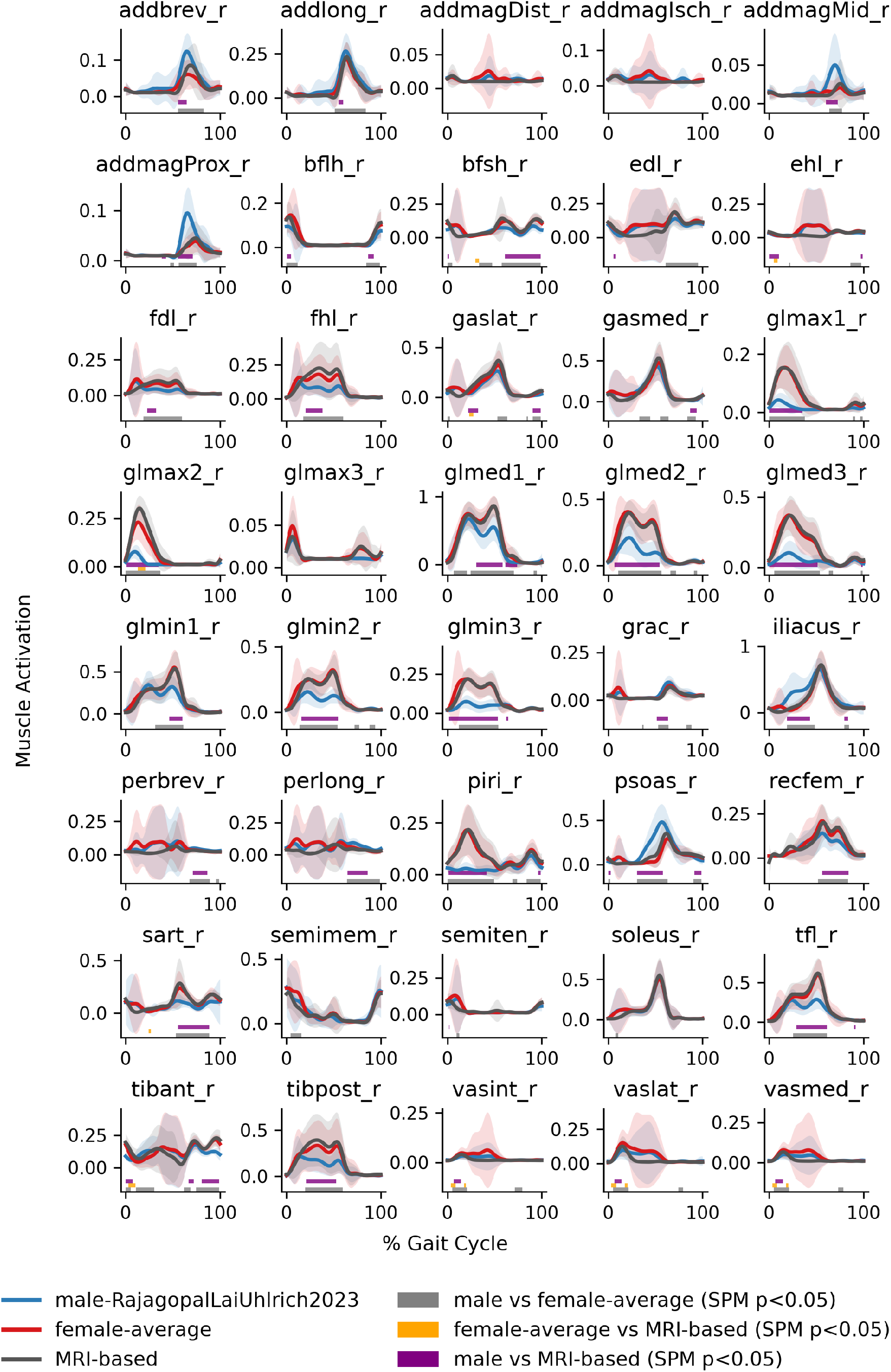
Waveforms and SPM analysis results for muscle activations (static optimisation), right foot cycle.

### Female Body Segment Inertial Parameters

#### Justification and Setup

In the MRI-based models, body segment inertial parameters (BSP; segment mass, centre of mass, and inertia tensor) were inherited from the linearly scaled gen-male model. The avg-female model therefore also retained these male-derived parameters. We tested whether replacing them with female-derived values affected the kinetic results.

We used the sex-specific data reported by de Leva (1996), which provide relative segment masses, centre-of-mass positions, and radii of gyration for a female sample (n = 15). Comparison with the model parameters showed that differences in segment mass were driven mainly by the underlying BSP convention rather than by sex (Table S5). For example, the model foot mass fraction was 61% higher than the de Leva female value and 52% higher than the male value. In contrast, the sex-specific component of these differences was relatively small (−5.8% to +11.1%).

**Table S5.**
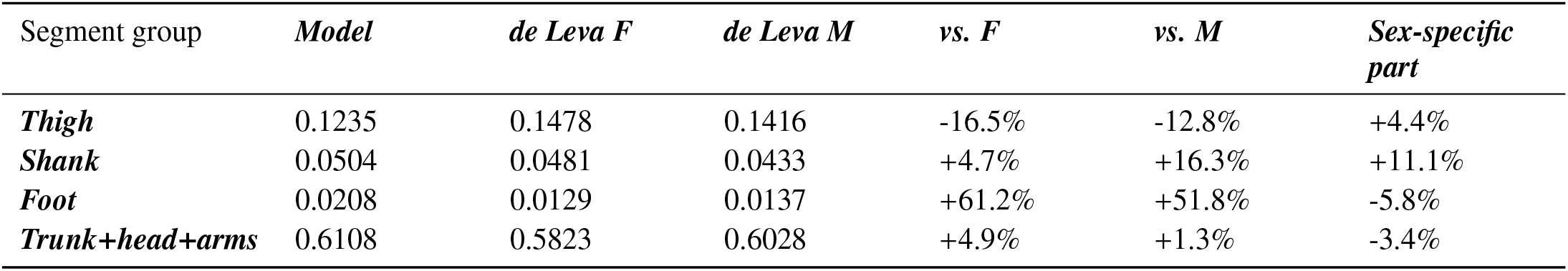
Model segment mass fractions (right side): direct comparison of model BSP against de Leva female values (right side).

Sex differences were more evident for centre-of-mass position and rotational inertia. The avg-female model placed the thigh centre of mass at 42.05% of segment length, compared with 36.12% in the de Leva female data and 40.95% in the male data. On the present femurs, the female–male difference in de Leva’s data corresponds to 19.0 mm. Within the de Leva dataset, female-to-male differences in rotational inertia were +31.3%, +23.4%, and +27.8% for thigh Ixx, Iyy, and Izz, respectively, and +25.7%, −9.6%, and +27.0% for the corresponding shank components. Shank centre-of-mass position differed by only 1.7 mm.

We therefore derived female BSP by applying the female-to-male magnitude and direction of the de Leva differences to the BSP of the avg-female model. The resulting multipliers are given in Table S6. Mass was adjusted first and then renormalised to conserve total body mass. Centre-of-mass position was adjusted only along the local longitudinal axis (the y-axis); the x- and z-components were unchanged.

**Table S6.** Final multipliers applied to each body (mass, realised mass after mass-conservation renormalisation, centre-of-mass y-component, and Ixx/Iyy/Izz excluding the mass ratio).

| *Body* | *Mass* | *Realised mass* | *CoM-y* | *Ixx* | *Iyy* | *Izz* |
| --- | --- | --- | --- | --- | --- | --- |
| **Femur** | 1.0438 | 1.0438 | 0.8821 | 1.2579 | 1.1821 | 1.2241 |
| **Tibia** | 1.1109 | 1.1109 | 0.9902 | 1.1316 | 0.8135 | 1.1430 |
| **Foot** | 0.9416 | 0.9416 | 0.9092 | 1.3536 | 1.2566 | 1.2968 |
| **Pelvis** | 1.1164 | 1.1164 | 1.0057 | 0.7746 | 0.8940 | 0.8317 |
| **Torso** | 0.9318 | 0.9170 | 0.9738 | 0.9170 | 0.8839 | 0.9324 |

For the thigh, shank, and foot, inertia multipliers were calculated as the squared ratios of the female-to-male radii of gyration, 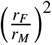. Segment-length ratios were not included because the model geometry was unchanged, and mass ratios were applied separately to avoid double-counting.

The pelvis and torso are not represented directly in the de Leva dataset. Their multipliers were therefore derived by combining the corresponding de Leva trunk components using parallel-axis composition. Sub-part lengths were normalised by sex-specific stature before calculating the ratios. This procedure required three approximations: no anterior–posterior offset was included in the Iyy transfer, products of inertia were left unchanged, and arm posture was assumed during torso assembly. Sensitivity testing across nine arm-posture combinations showed that posture had little effect on the resulting centre-of-mass, Ixx, and Izz ratios (0.31%, 1.23%, and 0.10%, respectively), although the effect on Iyy was larger (5.26%).

The final multipliers were applied to the mass, centre of mass, and inertia of each subject’s scaled and wrap-fixed avg-female model, producing a female-BSP variant of all 25 models while conserving total body mass. Because BSP do not affect inverse kinematics, the existing IK results were retained. Inverse dynamics, static optimisation, and joint reaction analyses were then repeated using the same processing pipeline. All 25 models were completed successfully.

Joint reaction forces were analysed separately for stance and swing using each subject-side’s recorded heel-strike and toe-off timings rather than a fixed percentage of the gait cycle. The mean stance duration was 63.0% (range 59.2–66.2%). Three subject-sides were excluded because the recorded cycle ended before toe-off, leaving 47 of 50 sides for the stance/swing analysis.

#### Results

Replacing the male-derived BSP with female-derived BSP had essentially no effect on the peak stance-phase hip and knee joint reaction forces (+0.05% and −0.06%, respectively; both p = 0.86; Table S7). The largest effect occurred at the back/lumbar joint, where peak force decreased by approximately 8.3% in both stance and swing. This reflects the 0.917 torso mass multiplier, which affects the lumbar joint consistently throughout the gait cycle. Although the corresponding test statistics were very large (|*t*| ≈ 90–94), this is a consequence of the deterministic nature of the simulations and should not be interpreted as a large practical effect. Smaller changes were observed at the ankle during stance (+0.08%) and the hip during swing (+2.23%). The remaining joint-phase combinations changed by less than 1.1%.

**Table S7.** Peak resultant joint reaction force (body weight normalised, paired comparison across 25 subjects, Holm-corrected across joint × phase).

| Joint | Phase | Male-derived BSP (BW) | Female BSP (BW) | Change | p (Holm) |
| --- | --- | --- | --- | --- | --- |
| Hip | Stance | 4.832 | 4.834 | +0.05% | 0.86 (n.s.) |
| Knee | Stance | 4.506 | 4.503 | -0.06% | 0.86 (n.s.) |
| Ankle | Stance | 6.250 | 6.255 | +0.08% | <0.001 |
| Back | Stance | 0.593 | 0.544 | -8.33% | <0.001 |
| Hip | Swing | 1.642 | 1.678 | +2.23% | 0.043 |
| Knee | Swing | 1.453 | 1.469 | +1.07% | 0.18 (n.s.) |
| Ankle | Swing | 1.013 | 1.009 | -0.43% | 0.86 (n.s.) |
| Back | Swing | 0.574 | 0.526 | -8.37% | <0.001 |

**Figure S5.**
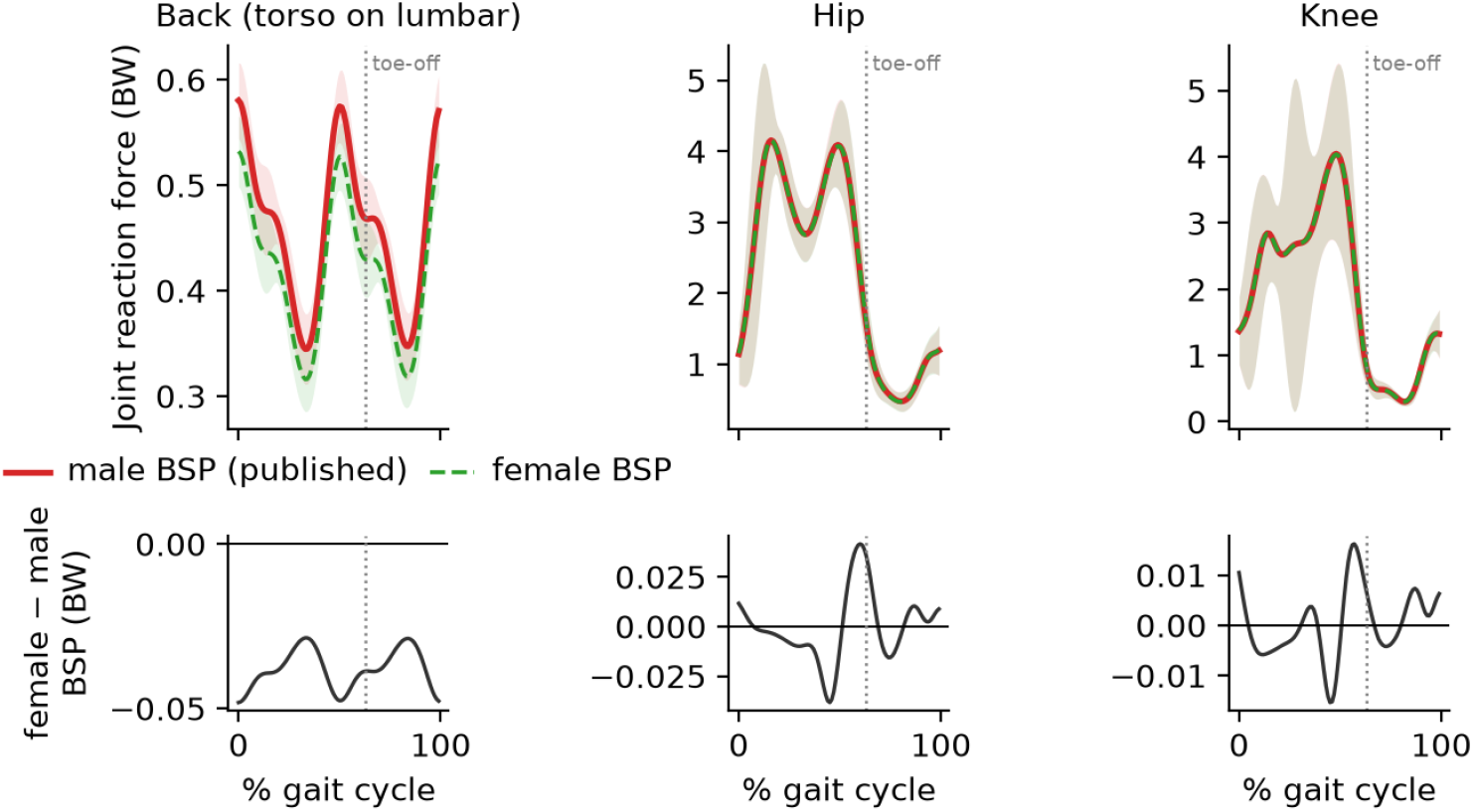
Waveforms for male- and female-derived BSP parameters and their effect on joint reaction forces.

#### Interpretation

Overall, replacing the male-derived BSP with female-derived values had negligible effect on the hip and knee joint reaction forces that are most relevant to the intended application. The main exception was the lumbar joint, where the lower torso mass produced an approximately 8% reduction in peak force. Thus, although female-specific inertial parameters produce statistically detectable changes at several joints, their practical effect on the primary lower-limb outputs is small.

Agreement with the MRI-based models changed only marginally (inverse kinematics was unchanged and inverse dynamics changed by 2.1%). However, these reference models also use male-derived BSP, so this comparison does not establish which inertial description is more accurate.

### Sensitivity to Inertial Parameters Study: Monte Carlo Propagation of Female BSP Uncertainty

#### Justification and Setup

To quantify the uncertainty introduced when applying the average-female model to individual women, inter-individual BSP variability was propagated through inverse dynamics using Monte Carlo simulation. de Leva (1996) reports mean values but not inter-individual standard deviations, so we used the DEXA-based study of 100 volunteers by Durkin and Dowling (2003), which reports within-group means and SDs by age and sex (Table 2). The Females 19–30 subgroup (n = 25; 57.0 kg, 163.5 cm, 22.0 years) was selected as the closest available match to the present cohort (59.6 kg, 165.6 cm, 26.2 years). Reported female coefficients of variation (SD/mean) were 6.23% for thigh mass, 5.15% for thigh centre of mass, and 3.54% for thigh radius of gyration (K); 6.56%, 1.90%, and 1.20%, respectively, for the leg; and 10.24% for foot mass.

The source has four limitations. First, it provides data for the forearm, hand, thigh, leg, and foot but not the trunk. Second, radius of gyration was reported only in the frontal plane, so applying it to all three model axes is an approximation. Third, foot centre of mass and radius of gyration were not reported because the original measurements had inconsistent axes in the supine position. Finally, the reported SDs include both biological variability and DEXA measurement error (validated to <3.2%). The smallest CVs, particularly the 1.2% for leg K, should therefore be regarded as upper bounds on the underlying biological variability rather than exact estimates.

Given these limitations, the sensitivity analysis perturbed the available limb-segment inertial properties using the SDs reported by Durkin and Dowling (2003), while holding trunk and foot centre of mass and radius of gyration fixed. Trunk mass was nevertheless allowed to vary through the subsequent renormalisation of segment masses to each subject’s measured total body mass. No separate trunk mass variability therefore had to be assumed. This setup provides an explicit lower-bound estimate of BSP uncertainty.

For each Monte Carlo draw, the subject’s scaled avg-female model was perturbed as follows. Segment mass was varied multiplicatively (*m* → *m*(1 + *e*)) and then renormalised so that total mass matched the subject’s measured mass. Thus, uncertainty was introduced only in the distribution of mass between segments. Centre of mass was perturbed along the segment’s longitudinal (y) axis only. Inertia was perturbed through the radius of gyration rather than directly through the inertia tensor:

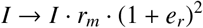

following *I* = *m*(*rL*)^2^, where *r*_*m*_ denotes the realised mass ratio. This avoids double counting the mass perturbation, consistent with the deterministic analysis above. Perturbations were constrained to maintain positive segment masses. For computational efficiency, only inverse dynamics was re-run for each draw. Because the preceding analysis showed that BSP changes do not affect inverse kinematics, each draw reused the existing IK solution and ground-reaction-force data. The uncertainties reported below therefore apply to joint moments only and were not propagated to the joint reaction forces reported in Table S7.

**Table S8.**
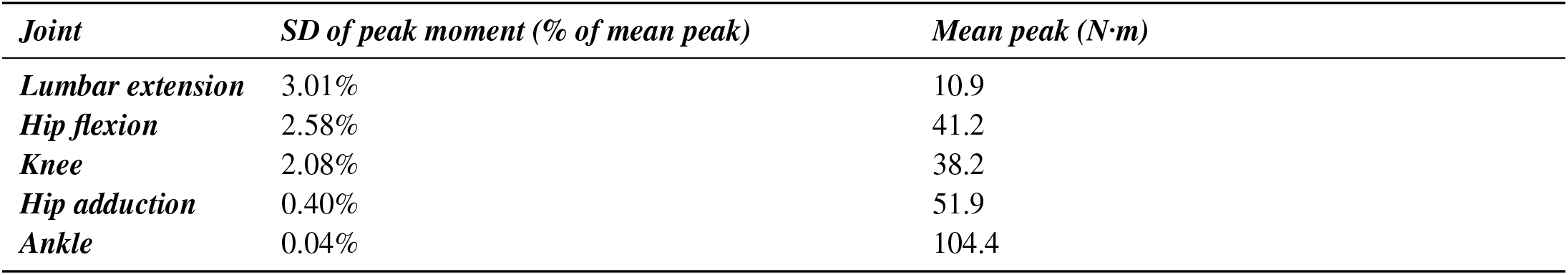
Uncertainty in peak joint moment under measured female inter-individual BSP variability.

#### Results

Applying the average-female model to another woman therefore introduces an estimated uncertainty of approximately 2–3% in peak hip-flexion and knee moments and essentially no uncertainty at the ankle. The lumbar joint shows the largest uncertainty. Because no measured female variability was available for the trunk, its centre of mass and radius of gyration were held fixed rather than assigned assumed values. The lumbar estimate is therefore a lower bound and is the result most likely to change if a trunk-inclusive source of female BSP variability becomes available.

#### Interpretation

The estimated inter-individual uncertainty associated with using population-average inertial parameters (2.1–2.6% for peak hip-flexion and knee moments) is substantially larger than the systematic difference between the male-derived and female-adjusted BSP at these primary outcomes. For hip and knee loading, therefore, the use of an average inertial description appears to matter more than which of the two average descriptions is used. The trunk is the exception. It is the segment for which no measured female variability was available and the source of the largest effect in the deterministic male-to-female BSP comparison (−8.3% in peak back joint reaction force). Applications of the model to trunk or lumbar loading should therefore treat the inertial description as an important source of uncertainty and should not rely on the present estimate without a trunk-inclusive source of female BSP variability.

